# The SARS-CoV-2 Nsp3 macrodomain reverses PARP9/DTX3L-dependent ADP-ribosylation induced by interferon signalling

**DOI:** 10.1101/2021.04.06.438552

**Authors:** Lillian Cristina Russo, Rebeka Tomasin, Isaac Araújo Matos, Antonio Carlos Manucci, Sven T. Sowa, Katie Dale, Keith W. Caldecott, Lari Lehtiö, Deborah Schechtman, Flavia Carla Meotti, Alexandre Bruni-Cardoso, Nicolas Carlos Hoch

**Affiliations:** Department of Biochemistry, Institute of Chemistry, University of São Paulo, São Paulo, Brazil; Faculty of Biochemistry and Molecular Medicine & Biocenter Oulu, University of Oulu, Oulu, Finland; Genome Damage and Stability Centre, School of Life Sciences, University of Sussex, Falmer, Brighton, United Kingdom; Current address: Department of Oncology, UCL Cancer Institute, University College London, London, United Kingdom

**Keywords:** COVID-19, SARS-CoV-2, ADP-ribosylation, PARP9, DTX3L, macrodomain

## Abstract

SARS-CoV-2 non-structural protein 3 (Nsp3) contains a macrodomain that is essential for virus replication and is thus an attractive target for drug development. This macrodomain is thought to counteract the host interferon (IFN) response, an important antiviral signalling cascade, via the removal of ADP-ribose modifications catalysed by host poly(ADP-ribose) polymerases (PARPs). Here, we show that activation of the IFN response induces ADP-ribosylation of host proteins and that ectopic expression of the SARS-CoV-2 Nsp3 macrodomain reverses this modification in human cells. We further demonstrate that this can be used to screen for cell-active macrodomain inhibitors without the requirement for BSL-3 facilities. This IFN-induced ADP-ribosylation is dependent on the PARP9/DTX3L heterodimer, but surprisingly the expression of Nsp3 macrodomain or PARP9/DTX3L deletion do not impair STAT1 phosphorylation or the induction of IFN-responsive genes. Our results suggest that PARP9/DTX3L-dependent ADP-ribosylation is a downstream effector of the host IFN response and that the cellular function of the SARS-CoV-2 Nsp3 macrodomain is to hydrolyse this end product of IFN signalling, and not to suppress the IFN response itself.

## Introduction

The SARS-CoV-2 pandemic has highlighted the need for an improved understanding of coronavirus pathogenesis for the development of novel antiviral strategies. The interferon (IFN) response is a central component of innate immunity and essentially precludes viral infection, as long as it is properly activated (Ivashkiv & Donlin, 2014; Schoggins, 2019). Therefore, successful replication of a virus within host cells requires active suppression or evasion of the host IFN response, which is often mediated by multiple viral proteins acting via separate mechanisms (Garcia-Sastre, 2017).

Type I IFN signalling is initiated upon recognition of viral nucleic acids in the cytoplasm, leading to the production and secretion of type I IFNs, such as IFNα and IFNβ, by virus-infected cells, whereas type II IFN, or IFNγ, is secreted by immune cells (Ivashkiv & Donlin, 2014; Lee & Ashkar, 2018). Binding of these cytokines to transmembrane IFN receptors induces activation of JAK kinases such as TYK2, JAK1 and JAK2, leading to phosphorylation and nuclear translocation of transcription factors of the STAT family, mainly STAT1, and subsequent induction of several hundred interferon-stimulated genes (ISGs) (Ivashkiv & Donlin, 2014; Schoggins, 2019). Among these ISGs are several members of the poly(ADP-ribose) polymerase (PARP) family, which catalyse the post-translational modification of proteins with ADP-ribose units using NAD^+^ as a substrate (Hoch & Polo, 2019).

ADP-ribosylation has recently emerged as a critical regulator of the IFN response, modulating central steps of this signalling cascade, both upstream of Type I IFN production and downstream of Type I or Type II IFN receptor binding (Fehr *et al*, 2020). Several IFN-regulated PARPs are also antiviral effectors, either by modifying host proteins involved in protein translation, stress granule formation and intracellular protein trafficking, or via modification and inhibition of viral proteins directly (Fehr *et al.*, 2020). Interestingly, some of these IFN-responsive PARPs, including PARP9, are rapidly evolving in the primate lineage, suggesting that they are engaged in an “arms race” with viral pathogens (Daugherty *et al*, 2014).

To counteract antiviral ADP-ribosylation by host PARPs, SARS-CoV-2 and other coronaviruses encode a macrodomain within non-structural protein 3 (Nsp3) that hydrolyses ADP-ribose modifications (Alhammad & Fehr, 2020; Alhammad *et al*, 2021; Leung *et al*, 2018). Importantly, inactivating mutations within this domain lead to reduced viral replication and increased activation of the host IFN response (Fehr *et al*, 2015; Fehr *et al*, 2016; Grunewald *et al*, 2019), strongly indicating that pharmacological inhibition of the Nsp3 macrodomain may be of substantial therapeutic value (Alhammad & Fehr, 2020; Brosey *et al*, 2021; Rack *et al*, 2020; Schuller *et al*, 2020).

Here, we show that activation of Type I or Type II IFN signalling induces ADP-ribosylation of host proteins that can be reversed by ectopic expression of the SARS-CoV-2 Nsp3 macrodomain in human cells. This ADP-ribosylation is dependent on the IFN-responsive PARP9/DTX3L heterodimer, but does not seem to modulate IFN signalling itself, since Nsp3 macrodomain expression or PARP9/DTX3L knockout had no effect on STAT1 phosphorylation or the induction of interferon-stimulated genes (ISGs). We propose that PARP9/DTX3L-dependent ADP-ribosylation of host proteins, which can be reversed by the SARS-CoV-2 Nsp3 macrodomain, is a downstream effector of the IFN response.

## Results

To study the functions of the SARS-CoV-2 Nsp3 macrodomain, we first engineered a sensitive assay to detect IFN-induced ADP-ribosylation in human cells. We transfected human A549 lung adenocarcinoma cells with the RNA mimetic poly(I:C), which induces a complete type I IFN cascade (Field *et al*, 1967; Matsumoto & Seya, 2008), or treated these cells with recombinant IFNγ, which induces type II IFN signalling (Kang *et al*, 2018; Wheelock, 1965). As expected, both treatments led to robust phosphorylation of STAT1 on tyrosine Y701 (Fig. 1A), which localized to the cell nucleus (Fig. 1B). Using the *Af1521* macrodomain-derived pan-ADP-ribose binding reagent (Millipore, MABE1016) for immunofluorescence staining and a high-content microscopy setup for robust image analysis and signal quantification (Sup.Fig. 1 and methods section), we observed a pronounced increase in a punctate, cytosolic ADP-ribose signal in response to either poly(I:C) or IFNγ treatment (Fig. 1C). Therefore, both type I and type II IFN signalling induce detectable ADP-ribosylation in human cells.

**Figure 1.**
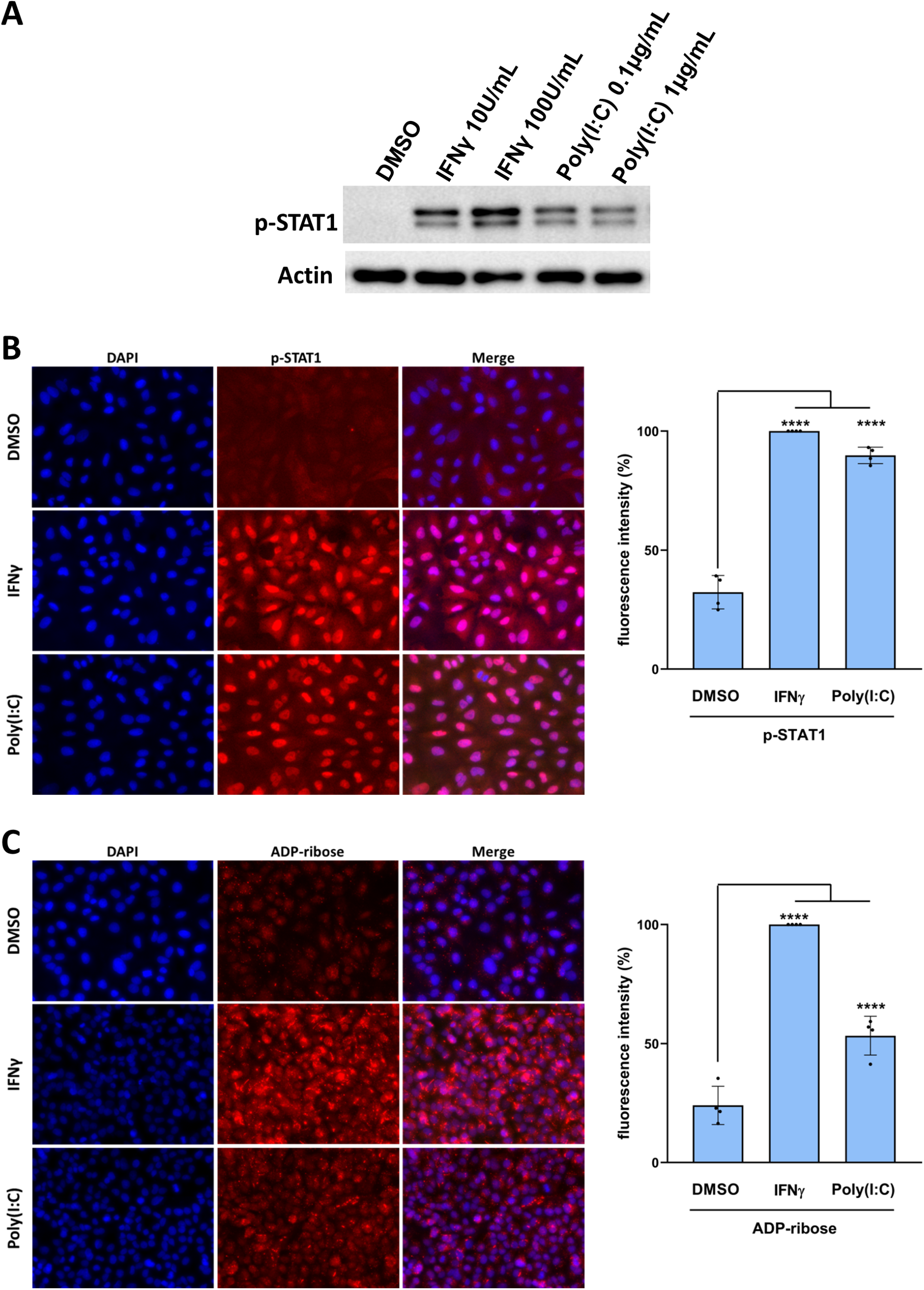
Both type I or type II IFN signalling induces ADP-ribosylation in A549 cells. **(A)** Immunoblot for STAT1 phospho-Y701 and actin loading control in A549 cells 24h after treatment with either vehicle control, recombinant interferon gamma (IFNγ) or transfection with poly(I:C), in the doses shown. **(B** (left) Representative images of immunofluorescence staining for STAT1 phospho-Y701 in A549 cells 24h after treatment with vehicle control, 100 U/mL IFNγ or transfection with 0.1 μg/mL poly(I:C); (right) quantification of mean p-STAT1 fluorescence per nucleus, averaged for thousands of cells per replicate and normalised to the IFNγ-treated sample. Mean ± SEM (n=4, from 3 separate experiments), ****= p<0.0001. **(C)** (left) Representative images of immunofluorescence staining for ADP-ribose modification (pan-ADP-ribose – Millipore) in A549 cells 24h after treatment with vehicle control, 100 U/mL IFNγ or transfection with 0.1 μg/ml poly(I:C); (right) Quantification of total ADP-ribose fluorescence in cytosolic dots per cell, averaged for thousands of cells per replicate.and normalised to the IFNγ-treated sample. Mean ± SD (n=4, from 3 separate experiments), ****= p<0.0001.

Next, we generated A549 cell lines ectopically expressing FLAG-tagged SARS-CoV-2 Nsp3 macrodomain using lentiviral vectors and confirmed that nearly 100% of the cells express the macrodomain using anti-FLAG immunofluorescence staining (Fig. 2A). Strikingly, constitutive macrodomain expression substantially reduced both poly(I:C) or IFNγ –induced ADP-ribosylation compared to empty vector controls (Fig. 2B). To confirm these results, we generated cells constitutively expressing a catalytically inactive N40A macrodomain mutant (Alhammad *et al.*, 2021; Fehr *et al.*, 2016) and repeated the experiment with the inclusion of recombinant IFNα and IFNβ. Again, expression of the WT macrodomain substantially reduced ADP-ribosylation induced by all of the treatments tested (Fig.2C). In contrast, expression of the N40A mutant had no appreciable effect on IFN-induced ADP-ribosylation (Fig. 2C), indicating that catalytic activity was required for this effect. In these experiments, we observed that the N40A mutant was expressed at lower levels than the WT macrodomain (Sup.Fig. 2A), which could, in principle, account for the higher persistence of ADP-ribose signal in the population of cells expressing the mutant protein. Therefore, the analysis in Fig. 2C includes only a subset of cells that express similar levels of WT and N40A macrodomain, based on anti-FLAG immunofluorescence intensity (Sup. Fig. 2B), which did not alter the original result (Fig. 2C and Sup.Fig. 2C-D). To further strengthen these observations, we generated A549 cell lines with doxycycline-inducible expression of either WT or N40A mutant macrodomain (Sup. Fig. 2E). As expected, IFNγ-induced ADP-ribosylation was indistinguishable between empty vector controls and either of the uninduced cells (Fig. 2D). In agreement with the results above, doxycycline-induced expression of the WT macrodomain reduced ADP-ribosylation, which was once again restored by the N40A mutation (Fig. 2D). As before, the cells used in this analysis were gated to ensure similar expression levels between WT and N40A mutant macrodomain, with no effect on the result (Sup.Fig.2F-G). Collectively, these data indicate that the Nsp3 macrodomain, when ectopically expressed in human cells, hydrolyses the ADP-ribosylation of host proteins induced by both type I or type II IFN signalling.

**Figure 2.**
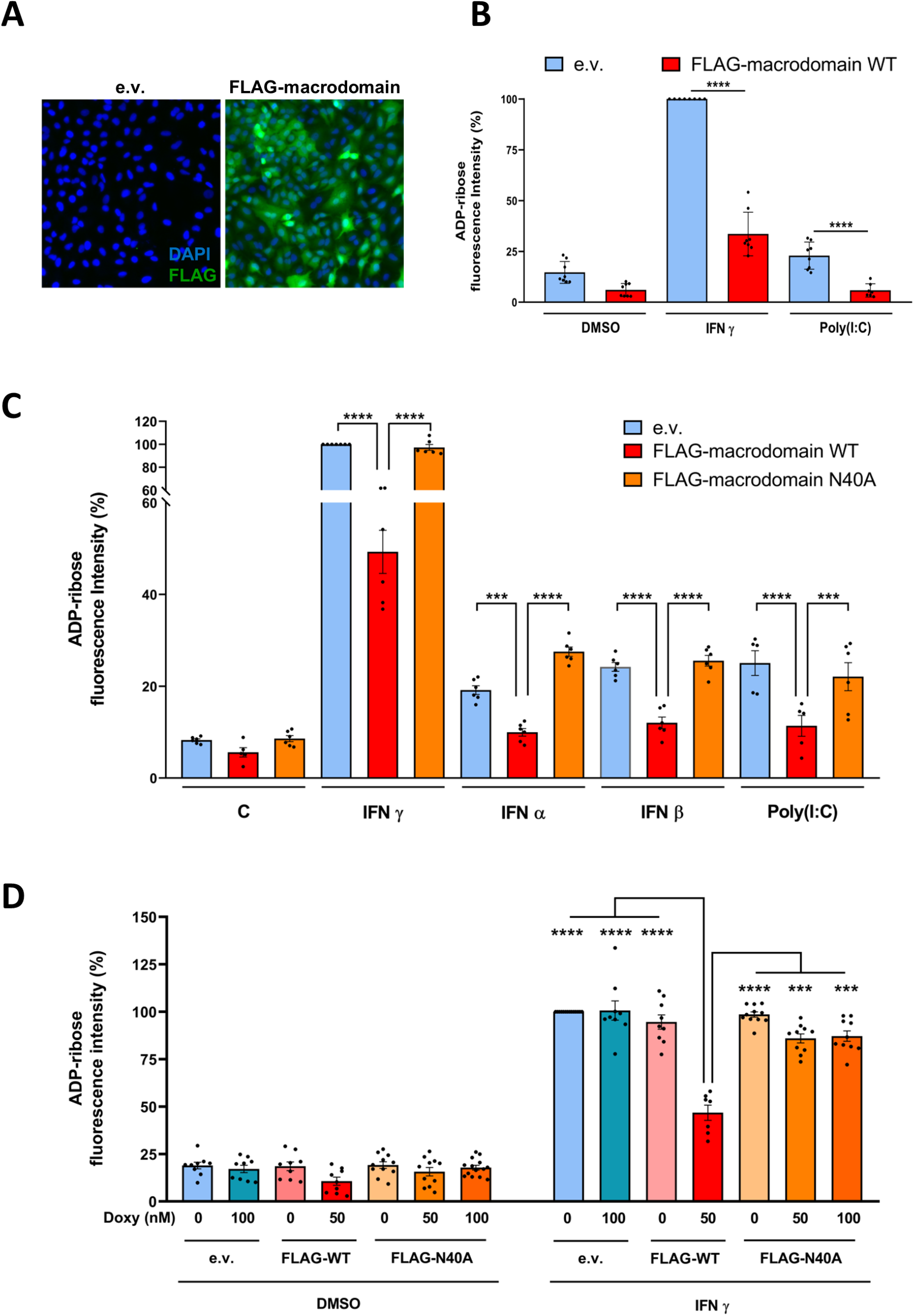
Ectopic expression of the Nsp3 macrodomain reverses IFN-induced ADP-ribosylation. **(A)** Representative images of anti-FLAG immunofluorescence staining (green) and DAPI staining (blue) in A549 cells transduced either with an empty vector control (left) or with a lentiviral construct constitutively expressing FLAG-tagged SARS-CoV-2 Nsp3 macrodomain (right). **(B)** Quantification of ADP-ribose immunofluorescence signal intensity in A549 cells transduced either with empty vector control (e.v.) or with a lentiviral construct for constitutive expression of FLAG-tagged macrodomain 24h after treatment with vehicle control, 100 U/mL IFNγ or transfection with 0.1 μg/ml poly(I:C). Mean ± SEM (n=8, from 3 separate experiments), ****= p<0.0001. **(C)** Quantification of ADP-ribose immunofluorescence signal intensity in A549 cells transduced either with empty vector control (e.v.) or with lentiviral constructs for constitutive expression of either WT macrodomain or catalytically dead N40A mutant, after 24h treatment with 1000 U/mL IFNα, 1000 U/mL IFNβ, 100 U/mL IFNγ, transfected with 0.1 μg/mL poly(I:C) or vehicle control. For FLAG-macrodomain-expressing samples, cells were gated such that macrodomain expression between WT and N40A mutant was comparable (Sup.Fig. 2A-C). Mean ± SEM (n=6, from 3 separate experiments), ****= p<0.0001, ***=p<0.001. **(D)** Quantification of ADP-ribose immunofluorescence signal intensity in A549 cells transduced either with empty vector control (e.v.) or with lentiviral constructs for doxycyclin-inducible expression of either WT macrodomain or catalytically dead N40A mutant, after 24h treatment with indicated doses of doxycycline and 100 U/mL IFNγ or vehicle control. For FLAG-macrodomain-expressing samples, cells were gated such that macrodomain expression between WT and N40A mutant was comparable (Sup. Fig. 2E-G). Mean ± SEM (n=7-12, from 4 separate experiments), ***= p<0.001 and ****= p<0.0001.

Given the urgent need for antiviral therapies for COVID-19 and the fact that the macrodomain is an attractive therapeutic target (Alhammad & Fehr, 2020; Brosey *et al.*, 2021; Rack *et al.*, 2020; Schuller *et al.*, 2020), we attempted to repurpose compounds that already have regulatory approval as potential Nsp3 macrodomain inhibitors. For this, we performed a structure-based virtual screen of a library of 6364 compounds that have been approved for human use by any regulatory agency in the world, against the deposited crystal structures of the SARS-CoV-2 Nsp3 macrodomain bound to ADP-ribose (PDB 6W02) (Fig. 3A and methods). Of the final list of 79 compounds of interest, 69 were sourced and tested in biochemical and cellular assays (Supplementary Table 1). Using thermal shift assays, which measure thermal stabilization of a protein upon ligand binding, we observed that, in contrast to the substantial shift in thermal stability of the Nsp3 macrodomain induced by the ADP-ribose positive control, none of the 69 test compounds showed any evidence of binding to the recombinant macrodomain (Fig. 3B). Similar results were obtained in cellular assays, in which none of the compounds substantially restored IFN-induced ADP-ribosylation levels in macrodomain-expressing cells (Fig. 3C), and the slight effect of atorvastatin observed in the cellular screen was not confirmed in follow-up experiments (Sup. Fig. 3).

**Figure 3.**
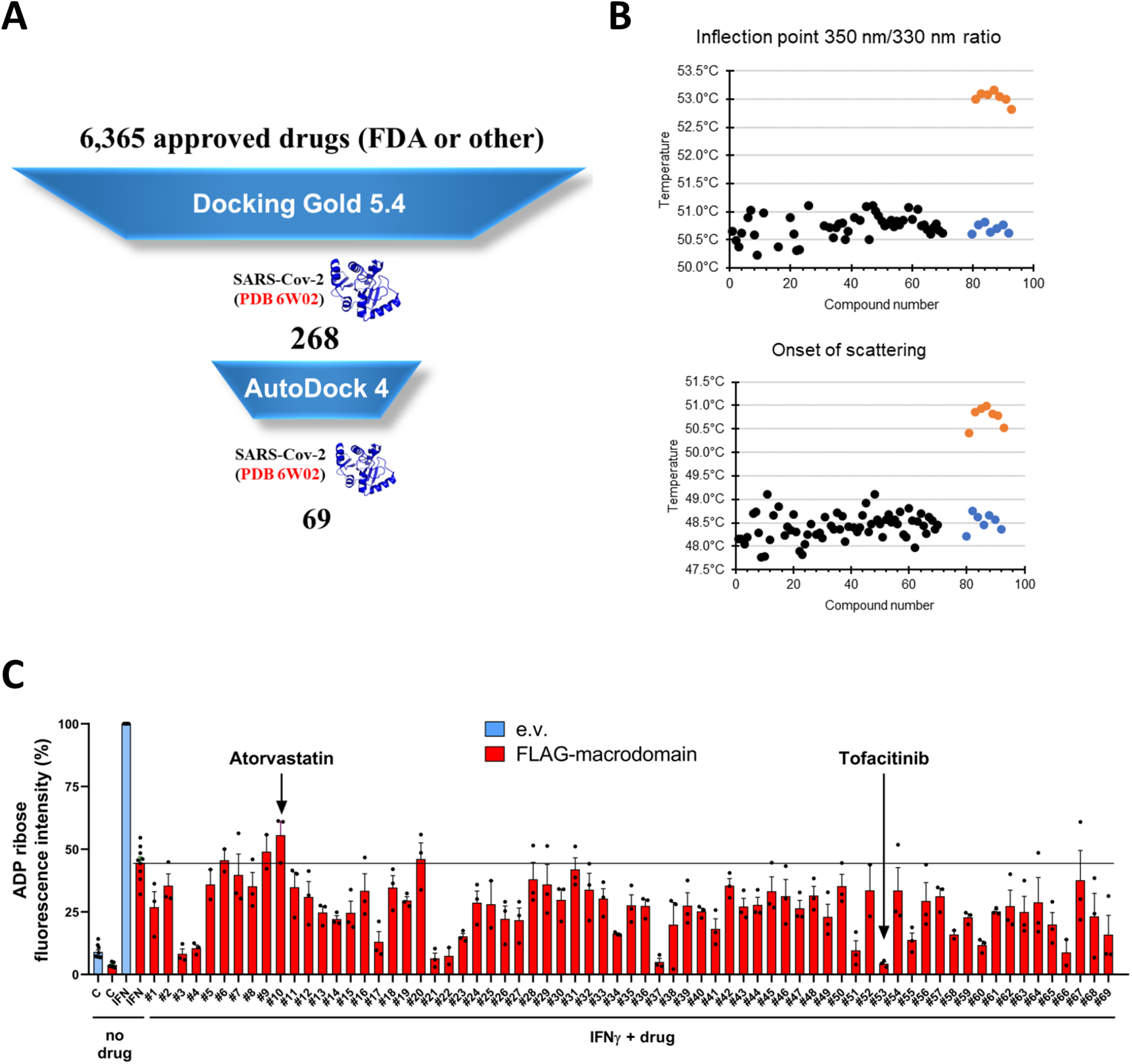
A repurposing screen for macrodomain inhibitors validates the technique for further screening. **(A)** Virtual screening workflow, starting with 6364 compounds approved for human consumption by any regulatory agency in the world, reaching a final list of 79 compounds taken forward for testing, of which 69 were sourced. **(B)** Thermal shift assays, by nanoDSF, of the recombinant Nsp3 macrodomain in the absence (blue) or presence of 100 μM ADP-ribose (orange) or 100 μM of each of the 69 test compounds (black). Melting temperatures measured as the inflection point of the 350 nm/330 nm intrinsic fluorescence ratio, indicative of protein unfolding (top) and onset of light scattering, indicative of protein aggregation (bottom) are shown. **(C)** Quantification of ADP-ribose immunofluorescence signal intensity in A549 cells transduced either with empty vector control (e.v.) or with a lentiviral construct for constitutive expression of WT macrodomain, 24h after treatment with vehicle control, 100 U/mL IFNγ or 100 U/mL IFNγ + 10-50 μM of each of 69 test compounds (Sup. Table 1). Mean ± SEM (n=3). Atorvastatin and tofacitinib (highlighted) are discussed in the main text.

Interestingly, we observed that some compounds reduced IFN-induced ADP-ribosylation (Fig.3C) and one of these was tofacitinib, an inhibitor of the JAK kinases that phosphorylate STAT1 in response to IFN receptor activation (Ghoreschi *et al*, 2011). As expected, tofacitinib treatment completely blocked STAT1-Y701 phosphorylation in response to IFNγ treatment (Fig. 4A, Sup.Fig. 4A) and this resulted in a complete loss of IFN-induced ADP-ribosylation (Fig. 4B, Sup.Fig. 4A). This suggests that ADP-ribosylation is likely to be catalysed by an IFN-responsive PARP that is transcriptionally induced by STAT1 complexes. This was corroborated by the fact that olaparib, an inhibitor of DNA-damage activated PARPs 1 and 2 that likely also inhibits PARP3, PARP4 and TNKS1/2 (PARP5a/b) at the relatively high concentration used here (Menear *et al*, 2008; Oplustil O’Connor *et al*, 2016; Thorsell *et al*, 2017), had no effect on IFN-induced ADP-ribosylation or on STAT1 phosphorylation (Fig. 4A-B, Sup.Fig. 4A).

**Figure 4.**
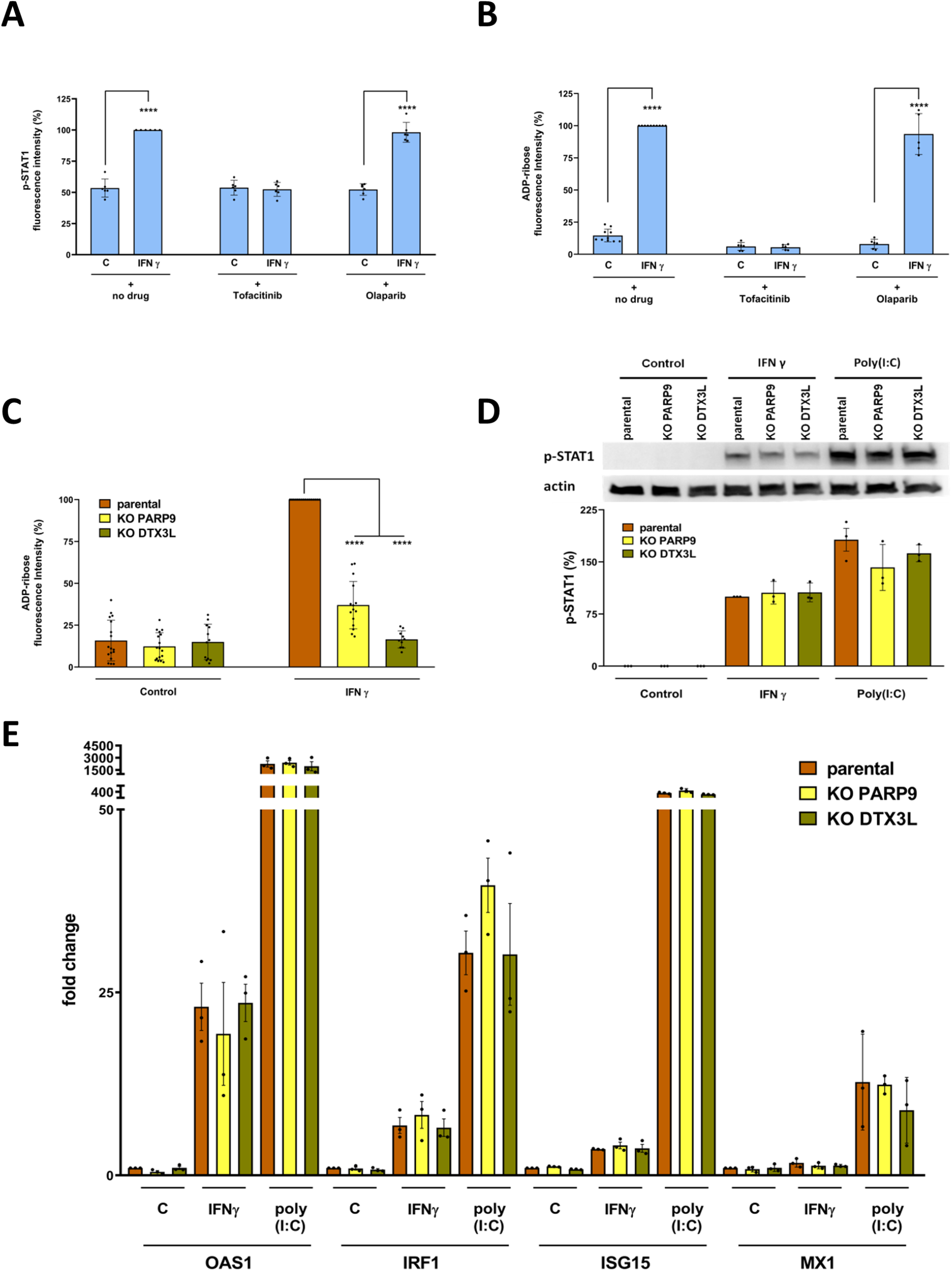
IFN-inducible PARP9 and DTX3L are required for IFN-induced ADP-ribosylation. **(A)** Quantification of nuclear STAT1 phospho-Y701 immunofluorescence signal intensity in A549 cells 24h after treatment with vehicle control or 100 U/mL IFNγ, with or without 10 μM tofacitinib or 10 μM olaparib, as indicated. Mean ± SEM (n=6, from 3 separate experiments), ****= p<0.0001. **(B)** Quantification of ADP-ribose immunofluorescence signal intensity (pan-ADP-ribose – Millipore) in A549 cells 24h after treatment with vehicle control or 100 U/mL IFNγ, with or without 10 μM tofacitinib or 10 μM olaparib, as indicated. Mean ± SEM (n=6-10, from 3 separate experiments), ****= p<0.0001. **(C)** Quantification of ADP-ribose immunofluorescence signal intensity (pan-ADP-ribose – Millipore) in WT, PARP9 or DTX3L KO RPE1-hTERT cells 24h after treatment with vehicle control or 100 U/mL IFNγ. Mean ± SEM (n=11-17, from 4 separate experiments), ****= p<0.0001. **(D)** Representative image (top) and quantification (bottom) of immunoblot analyses for STAT1 phospho-Y701 and actin loading control in WT, PARP9 KO or DTX3L KO RPE1-hTERT cells 24h after treatment with either vehicle control, 100U/mL IFN γ or transfection with 0.1 μg/mL poly(I:C). Mean ± SEM (n=3). **(E)** Relative levels of mRNA for OAS1, IRF1, ISG15 and Mx1 genes determined by RT-qPCR in WT, PARP9 KO or DTX3L KO RPE1-hTERT cells 24h after treatment with either vehicle control, 100U/mL IFNγ or transfection with 0.1 μg/mL poly(I:C), normalized to respective vehicle-treated WT cells. Mean ± SEM (n=3).

One such IFN-responsive PARP is PARP9, which forms a heterodimer with DTX3L and has been shown to participate in IFN-mediated antiviral responses (Juszczynski *et al*, 2006; Zhang *et al*, 2015). We had previously generated PARP9 and DTX3L CRISPR knockout cells in an RPE1-hTERT background (methods section and Sup. Fig. 4B-C) and decided to test whether IFN-induced ADP-ribosylation was altered in these cells. As with A549 cells, IFNγ treatment induced detectable ADP-ribosylation in RPE1 retinal pigment epithelial cells, confirming that this response is not specific to A549 cells (Fig.4C). Surprisingly however, PARP9 KO and particularly DTX3L KO cells displayed severely reduced IFNγ-induced ADP-ribosylation compared to controls (Fig. 4C, Sup.Fig. 4D). To determine if this was indirectly caused by an effect of PARP9 or DTX3L knockout on IFN signalling itself, we quantified STAT1-Y701 phosphorylation levels in response to IFNγ treatment in these cells, which revealed that PARP9 KO or DTX3L KO had no appreciable effect on STAT1 phosphorylation (Fig. 4D). To ascertain if the reduced ADP-ribosylation in PARP9 KO or DTX3L KO cells may be caused by a defect in IFN signalling downstream of STAT1 phosphorylation, we performed RT-qPCR to assess the transcriptional induction of four IFN-stimulated genes (ISGs)-OAS1, IRF1, ISG15 and Mx1 (Fujita *et al*, 1989; Haller *et al*, 1979; Reich *et al*, 1987; Shulman & Revel, 1980)-after treatment with either IFNγ or poly(I:C) (Fig. 4E). As expected, strong transcriptional induction of these ISGs was observed in RPE1 cells after IFNγ or poly(I:C) treatment, but knockout of PARP9 or DTX3L did not affect the induction of any of these genes under any of the conditions tested (Fig. 4E). These data indicate that PARP9 and DTX3L are essential for IFN-induced ADP-ribosylation of host proteins, which occurs downstream of ISG induction.

In light of these results, we decided to determine if Nsp3 macrodomain expression affects host IFN signalling or if its role is also downstream of ISG induction. Consistent with the latter, macrodomain-expressing A549 cells displayed normal levels of STAT1-Y701 phosphorylation in response to either IFNγ or poly(I:C) treatment (Fig. 5A, Sup.Fig. 5). Furthermore, expression of the Nsp3 macrodomain had no effect on the transcriptional induction of four ISGs in response to either IFNγ or poly(I:C) treatment (Fig. 5B). In agreement with this, macrodomain expression also did not impair the induction of PARP9 or DTX3L protein observed in response to IFNα, IFNβ, IFNγ or poly(I:C) treatment (Fig.5C). Therefore, these data indicate that the Nsp3 macrodomain does not affect the host IFN signalling cascade or the induction of interferon-stimulated genes, but is instead involved in suppressing downstream biological processes triggered by IFN-induced ADP-ribosylation (Fig. 6).

**Figure 5.**
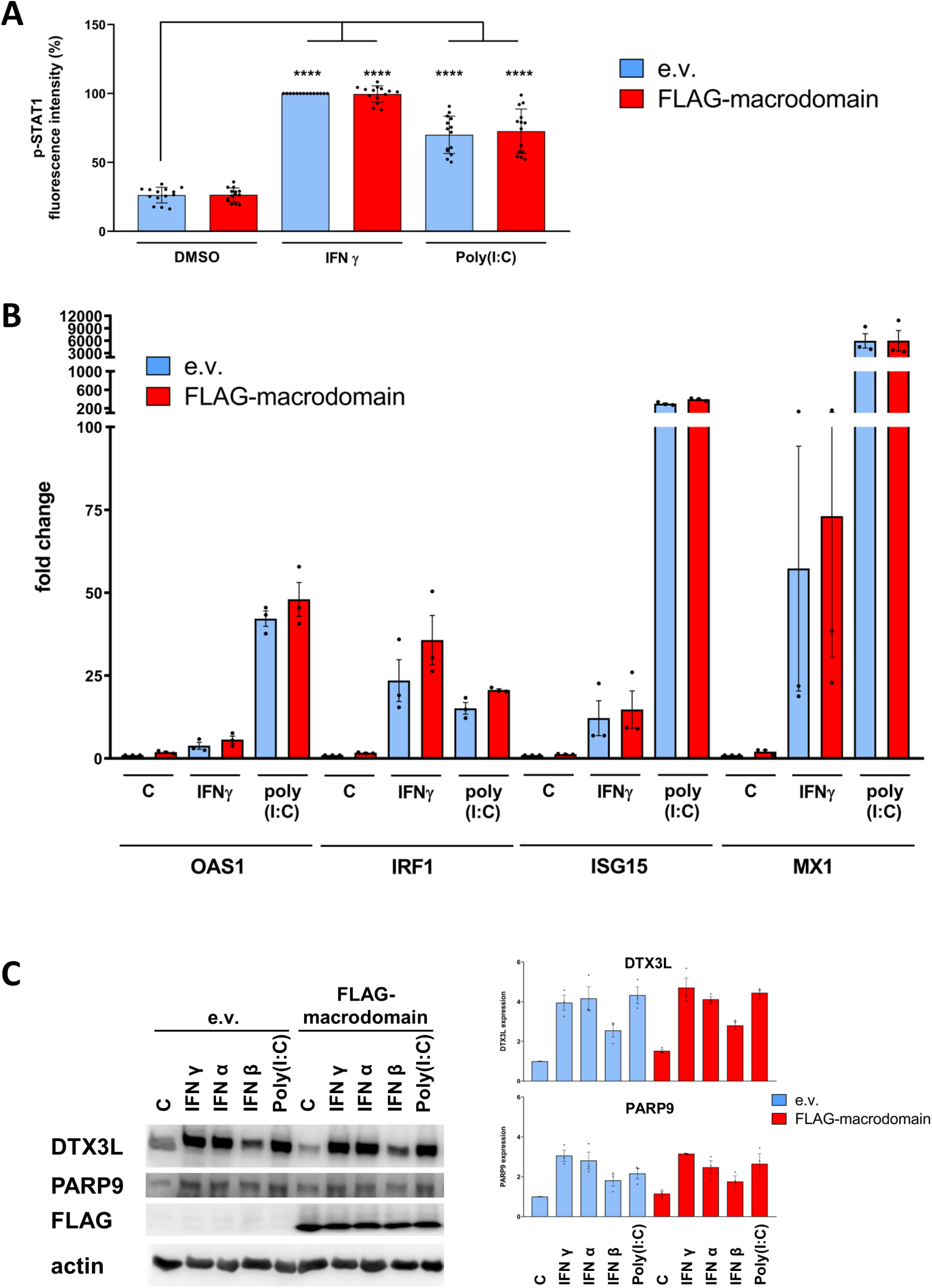
Nsp3 macrodomain expression has no effect on IFN signalling. **(A)** Quantification of nuclear STAT1 phospho-Y701 immunofluorescence signal intensity in A549 cells transduced either with empty vector control (e.v.) or with a lentiviral construct for constitutive expression of FLAG-tagged macrodomain, 24h after treatment with vehicle control, 100 U/mL IFNγ or transfection with 0.1 μg/ml poly(I:C). Mean ± SEM (n=14, from 3 separate experiments), ****= p<0.0001. **(B)** Relative levels of mRNA for OAS1, IRF1, ISG15 and Mx1 genes determined by RT-qPCR in A549 cells transduced either with empty vector control (e.v.) or with a lentiviral construct for constitutive expression of FLAG-tagged macrodomain, 24h after treatment with vehicle control, 100 U/mL IFNγ or transfection with 0.1 μg/ml poly(I:C), normalized to respective vehicle-treated empty vector control cells. Mean ± SEM (n=3). **(C)** Representative image (left) and quantification (right) of immunoblot analyses for DTX3L, PARP9, FLAG and actin loading control in A549 cells transduced either with empty vector control (e.v.) or with a lentiviral construct for constitutive expression of FLAG-tagged macrodomain, 24h after treatment with vehicle control, 1000 U/mL IFNα, 1000 U/mL IFNβ, 100 U/mL IFNγ or transfected with 0.1 μg/mL poly(I:C). Mean ± SEM (n=3).

**Figure 6.**
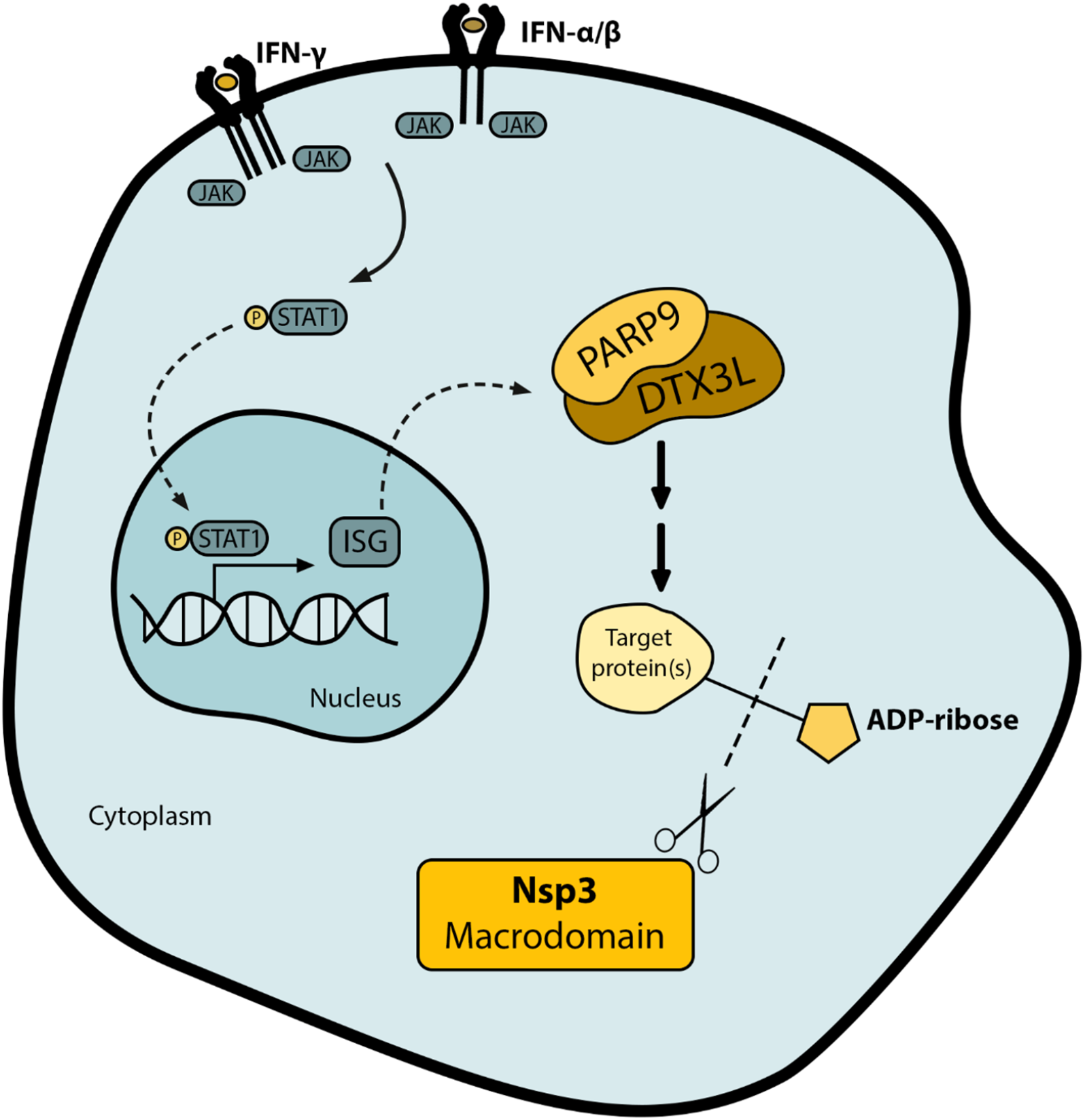
The macrodomain reverses PARP9/DTX3L-dependent ADP-ribosylation induced by IFN signalling. **(A)** Schematic representation of the proposed model. IFN signalling promotes STAT1 phosphorylation by JAK kinases and induces the expression of interferon-stimulated genes (ISGs), including PARP9 and DTX3L. This complex is required for downstream ADP-ribosylation of target proteins, which is counteracted by the viral Nsp3 macrodomain. Neither PARP9/DTX3L nor the Nsp3 macrodomain affect the IFN signalling cascade itself.

## Discussion

Successful viral evasion or suppression of the host interferon response is key for the establishment of a viral infection and is often mediated by several viral factors acting on multiple host targets (Garcia-Sastre, 2017). The coronavirus Nsp3 macrodomain is thought to represent an important mediator of coronavirus pathogenesis by reversing host antiviral ADP-ribosylation and is thus an attractive drug target (Alhammad & Fehr, 2020; Brosey *et al.*, 2021; Rack *et al.*, 2020; Schuller *et al.*, 2020). However, little is known about the molecular targets of IFN-responsive PARPs and how reversal of these modifications by the Nsp3 macrodomain may exert its pro-viral effects.

We have established a cellular immunofluorescence-based assay that can detect the physiological levels of ADP-ribosylation induced by both type I and type II IFN signalling in response to the viral RNA mimetic poly(I:C) or recombinant interferon treatments (Fig. 1 and 2), which is consistent with previous studies showing that IFNγ treatment results in detectable ADP-ribosylation during the process of macrophage activation (Higashi *et al*, 2019). We demonstrated that ectopically expressed SARS-CoV-2 Nsp3 macrodomain can hydrolyse these modifications in human cells (Fig. 2), which, to our knowledge, had so far only been shown *in vitro* using recombinant coronavirus Nsp3 macrodomains (Alhammad *et al.*, 2021; Fehr *et al.*, 2016).

We have employed this assay in a repurposing screen for Nsp3 macrodomain inhibitors (Fig.3), which turned out unsuccessful, but highlighted the reasonable throughput of our setup, which allowed two people to perform triplicate screens for 69 compounds in less than a week. Given the shortage of BSL-3 facilities required to test anti-coronavirus therapeutics on the SARS-CoV-2 virus directly, and the inherent uncertainty in these tests regarding on-target effects in case of successful reduction in viral loads, we believe that a cellular assay such as the one presented here may be a critical intermediate step in the development of Nsp3 macrodomain inhibitors, since a positive result in this assay ensures that candidate compounds are cell permeable and have on-target effects in human cells.

This microscopy-based assay also allowed us to investigate the origins and functions of the IFN-induced ADP-ribosylation. The signal we detected is predominantly cytosolic and has a punctate pattern (Fig. 1C) but, despite extensive efforts, we have so far been unsuccessful in co-localizing this signal with known cytosolic structures and organelles (LCR and NH, unpublished observations). Naturally, it will be critical to identify not only this structure but also the actual protein targets of the Nsp3 macrodomain-sensitive ADP-ribosylation induced by the IFN response, in order to elucidate the putative antiviral function of this modification.

Given that inhibition of DNA damage-responsive PARPs with olaparib had no effect on IFN-induced ADP-ribosylation, but inhibition of JAK kinases completely prevented its formation (Fig. 4A and B), our data indicate that the ADP-ribosylation detected here is catalysed by a PARP whose expression is induced by IFN signalling. Importantly, we identified that the PARP9/DTX3L heterodimer, whose expression is IFN-stimulated (Fig. 5C), is absolutely necessary for this ADP-ribosylation, as the signal was lost upon knockout of either of these genes (Fig. 4). Interestingly, PARP9 and DTX3L expression is also induced in cells infected with SARS-CoV-2, as long as viral load is high enough to induce an IFN response (Blanco-Melo *et al*, 2020). In agreement with our results, PARP9/DTX3L induction under these conditions is completely prevented by treatment with a JAK kinase inhibitor (Blanco-Melo *et al.*, 2020). While it is tempting to speculate that PARP9/DTX3L directly catalyse the formation of the IFN-induced ADP-ribose signal we detected, we cannot rule out a role of these proteins in a putative signalling cascade that culminates in ADP-ribosylation by another PARP.

We propose that the PARP9/DTX3L heterodimer promotes this ADP-ribosylation as a downstream effector of the IFN response, as we did not observe an effect of PARP9/DTX3L deletion on STAT1 phosphorylation or ISG induction (Fig. 4). These data are surprising, considering previous reports showing that PARP9 and DTX3L overexpression increases ISG induction in response to IFN signalling in human fibrosarcoma cells (Zhang *et al.*, 2015) and that PARP9 silencing reduces IFNγ-induced STAT1 phosphorylation and ISG induction in macrophages (Iwata *et al*, 2016). While further studies are clearly required to clarify the role of PARP9 and DTX3L in the IFN response, we speculate that differences in genetic manipulation of PARP9/DTX3L levels (overexpression, RNA interference or CRISPR/Cas9 KO) or cell types (fibrosarcoma, macrophage or retinal pigment epithelia) could account for these differences.

In agreement with our model for an effector function of this PARP9/DTX3L–dependent ADP-ribosylation, the ectopic expression of the Nsp3 macrodomain also had no effect on STAT1 phosphorylation or ISG induction (Fig. 5), despite reversing PARP9/DTX3L-mediated ADP-ribosylation (Fig. 2). This was again surprising, since macrodomain mutation in a mouse coronavirus model (mouse hepatitis virus- MHV) was previously shown to cause an increase in IFN production by virus-infected murine macrophages (Grunewald *et al.*, 2019). In this MHV model, the macrodomain was proposed to mainly counteract PARP14 activity (Grunewald *et al.*, 2019), which is thought to act upstream of the production of type I IFN by virus-infected cells (Caprara *et al*, 2018). These results are not necessarily conflicting, as the MHV macrodomain may hydrolyse different PARP targets that play a more prominent role in the mouse IFN response relative to humans (Zschaler *et al*, 2014). Alternatively, macrodomain mutation in the context of a full length Nsp3 protein may have secondary effects on the activity of other domains, such as the papain-like protease known to suppress IFN signalling (Li *et al*, 2021; Shin *et al*, 2020).

In conclusion, we show here that the SARS-CoV-2 Nsp3 macrodomain hydrolyses PARP9/DTX3L-dependent ADP-ribosylation induced by IFN signalling and uncover a role for this modification as a putative effector, rather than modulator, of the IFN response (Fig. 6). As part of this study, we developed a cellular assay with the potential to substantially impact drug discovery efforts currently underway to target the Nsp3 macrodomain as a novel anti-coronavirus therapy.

## Materials and Methods

### Cell culture and treatment conditions

A549 lung adenocarcinoma cells were grown in DMEM/high glucose media (Thermo) and RPE1-hTERT retinal pigment epithelia cells were grown in DMEM/F-12, both supplemented with 10% Fetal Bovine Serum (Thermo). HEK293 FT cells were grown in DMEM/high glucose supplemented with 10% FBS (Thermo), 50mg/mL gentamycin (Sigma), 1mM sodium pyruvate (Thermo), non essential amino acids (Thermo) and 2mM L-glutamine (Thermo). All cell lines were maintained at 37°C in a humidified atmosphere containing 5% CO_2_. Recombinant interferons α, β and γ (Sigma SRP4594, I9032 and SRP3058) were added directly to the media at the indicated doses and poly(I:C) (Sigma) was transfected using PEI and the cells collected 24h later. Tofacitinib (Selleckchem), olaparib (Selleckchem) or atorvastatin (Sigma) were added at the indicated doses at the same time as the induction of IFN signalling.

### Lentiviral construct generation and viral transduction

SARS-CoV2-Nsp3 macrodomain (UniProt identifier P0DTC1, residues 1023 to 1157) was amplified from a pET30a-based vector (a kind gift by A.Fehr - Univ. of Kansas), and cloned via BamHI and XhoI sites into a pCDNA3.1-based vector previously engineered to contain an N-terminal FLAG-FLAG-Strep-Strep tag. This ORF was then subcloned into a pCDH-puro lentiviral vector (System Biosciences) via NheI and XhoI sites or amplified and cloned via EcoRI and AgeI sites into the pLVX-puro lentiviral vector (System Biosciences). The N40A mutation was generated by standard site-directed mutagenesis and all vectors were confirmed by Sanger sequencing of the inserts. These vectors were co-transfected with psPAX2 and pMD2.G packaging vectors (Systems Biosciences) into HEK-293FT cells using standard PEI transfection, the supernatant collected 48h and 72h after transfection, filtered through 0.45 μm filters and used to transduce A549 cells in the presence of 8 μg/mL polybrene (Sigma). After 48 hours, 5 μg/mL puromycin was added to the media for selection of transduced cells for 7 days.

### Virtual Screening

The crystal structure of the Nsp3 macrodomain from SARS-CoV-2 co-crystalysed with ADP-ribose (Protein Data Bank code 6W02) was used for structure-based studies (Michalska *et al*, 2020). The protonation states of residues were revised and the variable conformation of the Asp27 residue was adjusted. Hydrogen atoms were added and water molecules and ligands removed. Molecular docking simulations were performed with Gold 5.4 and AutoDock 4.2.3 softwares. The protein was set as rigid and the ligands as flexible. For the docking studies using Gold 5.4, the radius of simulation was set to 6 Å (considering the large size of ADP-ribose). CHEMPLP score function was selected and the efficiency parameter set to very flexible, while other parameters were left at default values. For the docking with AutoDock, non-polar hydrogen atoms were merged with respective carbons and Gasteiger charges were added. The number of points of each dimension was 25 × 40 × 50 Å and grid box was centralized in the coordinates 10.649 × 7.101 × 20.841. For each ligand, 100 simulations of molecular docking were performed. For optimization, Lamarckian genetic algorithms were employed and the molecular docking protocols were validated by redocking. For this, co-crystallized ADP-ribose was removed and re-docked in PDB 6W02. RMSD distances between experimental and simulated atom ligands were calculated by VMD (Humphrey *et al*, 1996). For virtual screening, the subset of 6364 drugs approved in the world available in Zinc15 (http://zinc15.docking.org/) was select as target library.

### Cloning of expression vectors and purification of recombinant protein

The SARS-CoV-2 Nsp3 macrodomain (Uniprot identifier P0DTC1, residues 1024-1192) was cloned into pNH-TrxT vector (Addgene plasmid #26106) using SLIC restriction free cloning method (Jeong *et al*, 2012). Briefly, the pNH-TrxT plasmid was linearized and 100 ng of linearized plasmid were mixed in a 1:4 molar ratio with SARS-CoV-2 Nsp3 macrodomain (gBlock gene fragment, Intergrated DNA Technologies) and incubated with T4 DNA polymerase for 2.5 min at room temperature and for 10 min on ice. The mixture was used to transform NEB5α competent *E. coli* cells (New England BioLabs) according to manufacturer’s instructions. Colonies were grown on LB agar containing 5% sucrose using the SacB-based negative selection marker (Hynes *et al*, 1989). The construct was verified by sequencing of the insert regions, and was then used to transform *E. coli* BL21(DE3) cells. 500 mL Terrific Broth (TB) autoinduction media including trace elements (Formedium, Hunstanton, Norfolk, England), supplemented with 8 g/l glycerol and 50 μg/ml kanamycin, was inoculated with 5 mL of overnight preculture and incubated at 37°C until an OD600 of 1 was reached. After an overnight incubation at 16°C, the cells were collected by centrifugation at 4,200×g for 30 min at 4°C.

The pellets were resuspended in lysis buffer (50 mM HEPES pH 7.5, 500 mM NaCl, 10 mM imidazole, 10% glycerol, 0.5 mM TCEP) and stored at − 20°C. For protein purification, the cells were thawed and lysed by sonication. The lysate was centrifuged (16,000×g, 4°C, 30 min), filtered and loaded onto a 5 ml HiTrap HP column equilibrated with lysis buffer and charged with Ni^2+^. The column was washed with 30 column volumes of lysis buffer and 4 column volumes of wash buffer (30 mM HEPES pH 7.5, 30 mM imidazole, 500 mM NaCl, 10% glycerol, 0,5 mM TCEP). The protein was eluted using wash buffer containing 300 mM imidazole. Imidazole was removed by dialysis and TEV-protease was added (1:30 molar ratio, 16 h, 4°C) to cleave the His_6_-TrxT-tag followed by a reverse IMAC step to remove impurities. Size exclusion chromatography was carried out on a HiLoad 16/600 Superdex 75 pg 120 mL column in 30 mM HEPES pH 7.5, 300 mM NaCl, 10% glycerol and 0,5 mM TCEP. Pure fractions were pooled and stored at −70°C.

### Thermal shift assay

The purified SARS-CoV-2 Nsp3 macrodomain was diluted to 0.3 mg/ml in assay buffer (25 mM HEPES pH 7.5, 100 mM NaCl) and mixed with 100 μM of the compounds. The sample containing Flubendazole was measured at 20 μM compound concentration due to limited solubility. Samples in the presence or absence of 100 μM ADP-ribose were used as controls. All samples and controls contained a final concentration of 1%(v/v) DMSO. Samples were loaded to glass capillaries and analysis was performed in Prometheus NT.48 (NanoTemper). Data points were recorded from 35– 65°C, with the temperature increasing by 1°C/min. The onset of scattering and melting temperatures based on the change of intrinsic protein fluorescence (ratio 350 nm/330 nm) were calculated in PR.ThermControl software (NanoTemper).

### CRISPR/Cas9 knockout generation

PARP9 and DTX3L sgRNA sequences were 5’ GATCTGATGGGATTCAACG (exon 8) and 5’GCAGTTCGCTGTATTCCA (exon 4), respectively. Appropriate oligonucleotides were annealed, phosphorylated and cloned into BbsI-digested eSpCas9(1.1) vector (Addgene #71814). We subsequently discovered that these gRNAs, which were designed as tru-gRNAs with 17bp of homology plus a 5’G (Fu *et al*, 2014) are incompatible with the eSpCas9(1.1) mutant (Slaymaker *et al*, 2016). Therefore, RPE1-hTERT cells were co-transfected with either of these vectors and the hCas9 vector (Addgene #41815) using a Neon Transfection system (Thermo) and transfected cells selected with G418 for 5 days. Individual clones were screened by western blotting and clones with complete absence of PARP9 and DTX3L protein were selected (Sup. Fig. 4B). Genomic DNA surrounding the edited locus was amplified and analysed by Sanger sequencing (Sup.Fig. 4C).

### Western blotting

Adherent cells were washed in PBS and lysed directly in pre-heated Laemmli buffer devoid of bromophenol blue and beta-mercaptoethanol. Lysates were transferred to tubes, boiled for 15min and the protein concentration determined using BCA protein quantification kit (Pierce). After normalization of protein concentrations and addition of bromophenol blue and beta-mercaptoethanol, samples were boiled again for 10min and 15-50 μg of protein were loaded per sample in standard SDS-PAGE gels. Proteins were transferred to nitrocellulose membranes (Bio-Rad), visualized with Ponceau Red (Sigma) and the membranes cut horizontally such that different portions of the same membrane could be incubated with the appropriate antibodies. Membranes were blocked with 5% BSA for 30 min and incubated with primary antibody overnight at 4°C. After extensive washing in TBST buffer, 1h incubation in appropriate HRP-conjugated secondary antibodies (Sigma) and another round of washing, membranes were incubated with ECL Prime (Amersham) and the signal detected using a Chemidoc MP Imaging System (Bio-Rad). Signals were quantified using ImageJ software.

### Immunofluorescence staining

Cells were seeded either on 1.5H glass coverslips (Thorlabs) or in microscopy-compatible plastic 96-well plates (Corning), treated as required, washed with PBS and fixed with 2% (for pSTAT1) or 4% EM-grade PFA (EMS) prepared in PBS, which was subsequently quenched with 0.1M glycine. After permeabilization in 100% methanol (for pSTAT1) or 0.2% TritonX-100 in PBS, samples were blocked in 1% BSA/5% goat serum in PBS and incubated with primary antibody for 1h at room temperature (for ADP-ribose) or overnight at 4°C. Samples were extensively washed in PBS, incubated with appropriate fluorescently labelled secondary antibodies (Thermo), washed again, stained with DAPI (Thermo) and the coverslips mounted in Vectashield (Vector Labs) or the plates maintained in 90% glycerol until image acquisition.

### Image acquisition and analysis

Fluorescence microscopy images were acquired on a customized TissueFAXS i-Fluo system (TissueGnostics) mounted on a Zeiss AxioObserver 7 microscope (Zeiss), using 20x Plan-Neofluar (NA 0.5) or 40x Plan-Apochromat (NA 0.95) objectives and an ORCA Flash 4.0 v3 camera (Hamamatsu). For most experiments, 6×6 adjacent fields of view were acquired per sample using automated autofocus and image acquisition settings. Images were analysed using StrataQuest software (TissueGnostics). For phospho-STAT1 quantification, individual nuclei were detected in the DAPI channel and the mean intensity of phospho-STAT1 signal per nucleus quantified for thousands of cells per sample. For ADP-ribose quantification, individual nuclei were detected in the DAPI channel and the approximate cell boundaries determined by growing outwards from the nuclear area for a fixed 30 microns, until neighbouring areas touched each other or until the ADP ribose signal reached background levels. Peaks of ADP-ribose signal intensity within this cellular mask (excluding the nucleus) were detected as “dots” and the total fluorescence signal contained in all dots per cell was quantified for thousands of cells per sample. In both cases, the fluorescence intensities of all cells within a sample were averaged and this value normalized to the average intensity of the IFNγ-treated control sample for each biological replicate experiment (Sup.Fig. 1).

### RT-qPCR

Total RNA was extracted from 10^6^ cells using an RNAeasy kit (Qiagen), treated with DNAseI (Ambion) and reverse-transcribed using SuperScriptII (Thermo), according to manufacturer instructions, using both oligodT and random hexamer primers (Thermo). This cDNA was used for qPCR (5 ng/reaction) using Power SYBR green Master mix (Thermo) with three technical replicates per biological replicate, using 200 nM of the primer sets indicated in Table 1, which were chosen from PrimerBank (Wang *et al*, 2012). Reactions were performed on an Applied Biosystems 7500 Real-time PCR system, using default settings. RPL19 was used as housekeeping control and standard 2^−ΔΔCt^ analysis performed relative to the untreated control sample.

**Table 1.**
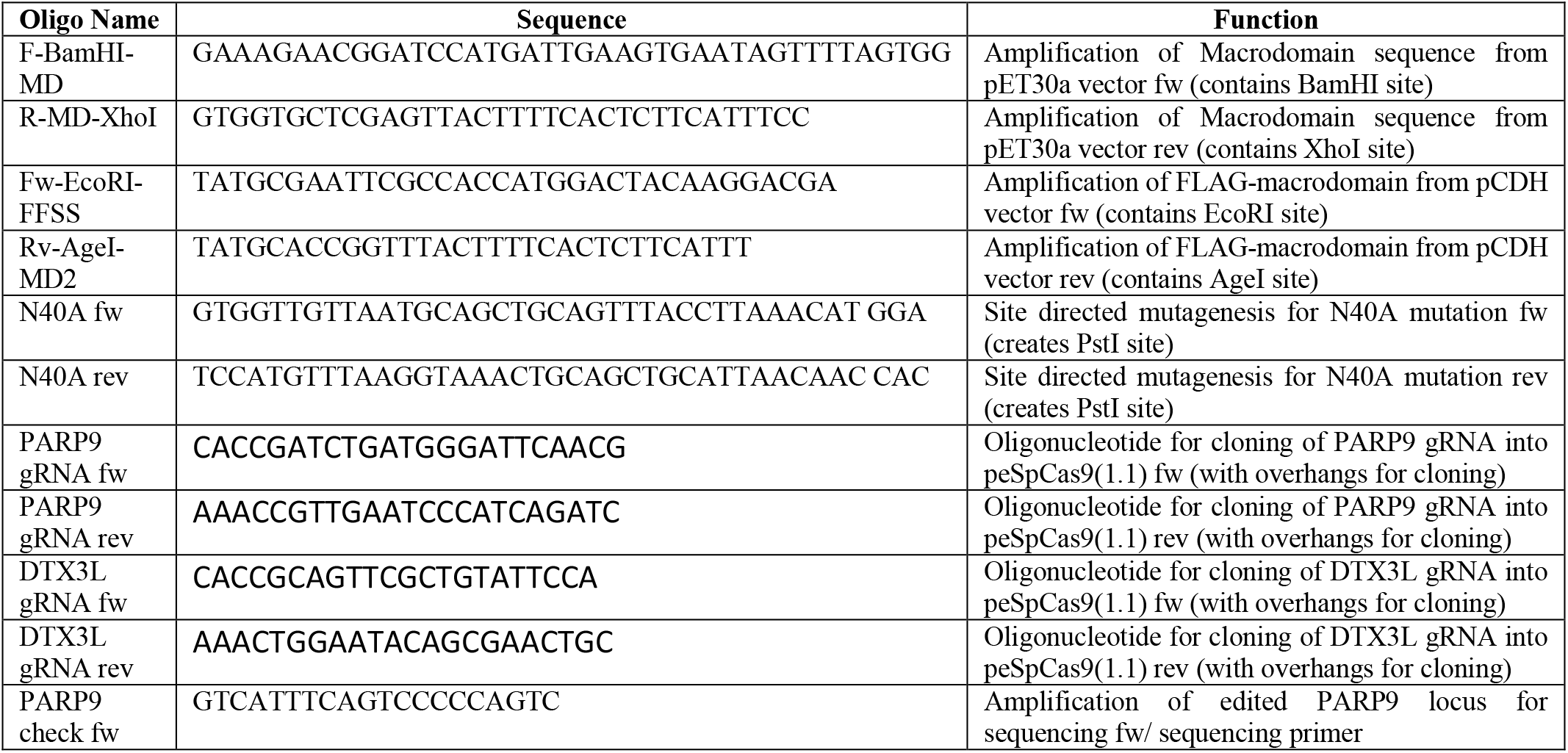
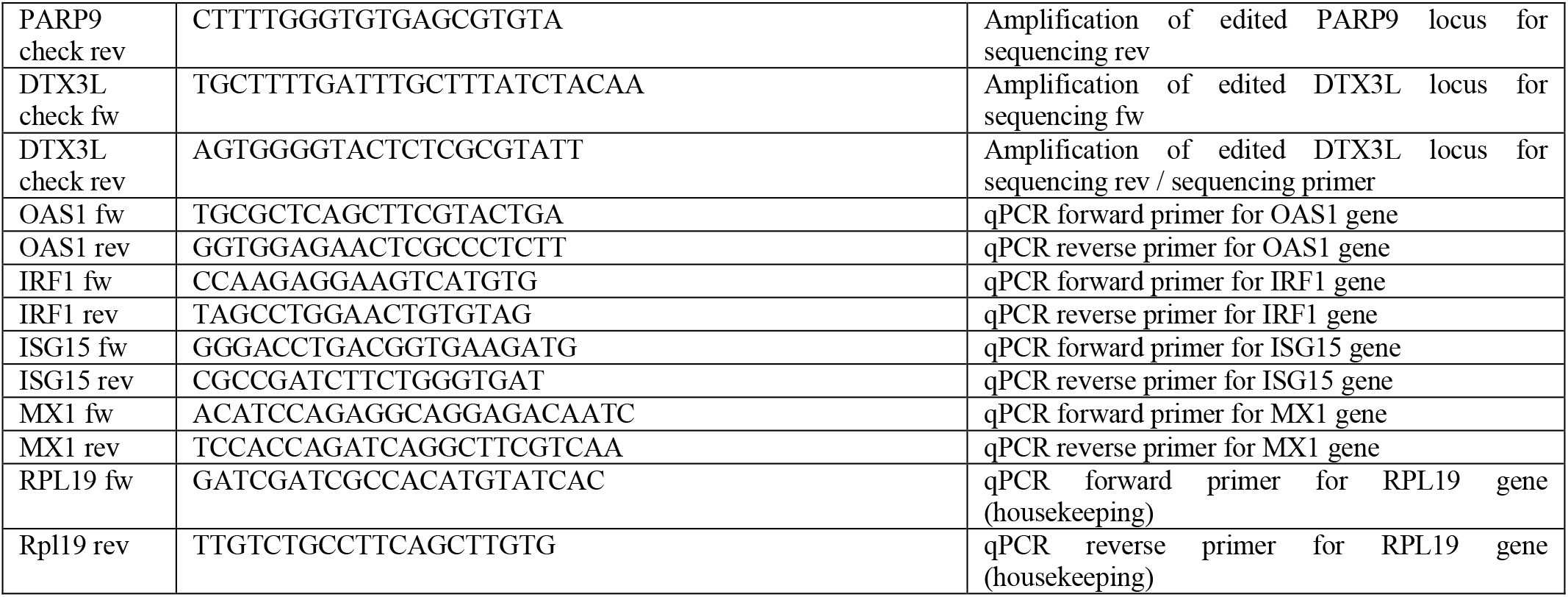
Oligonucleotides sequences and their functions:

### Statistical analyses

All experiments were repeated on at least three separate occasions, often with multiple parallel replicates processed on the same day, but treated as independently as possible. All graphs and statistical analyses were generated using GraphPad Prism software and display mean ± SEM of the normalized values relative to the IFNγ-treated control for each replicate, considered as 100%. Statistical comparisons between samples were performed using ANOVA, with p<0.0001 indicated by ****, p<0.001 indicated by *** and p<0.01 indicated by **.

### Primers and Antibodies

All oligonucleotides and antibodies used in this study are presented in the tables below:

**Table 2.** Antibodies and their respective applications

| Antibody | Host Species | Supplier (Catalogue Number) | Application (Dilution) |
| --- | --- | --- | --- |
| Actin | Mouse | Millipore (MAB1501) | WB (1:5000) |
| pan-ADP-ribose | Rabbit | Millipore (MABE1016) | IF (1:400) |
| pan-ADP-ribose | Mouse | A kind gift by M. Hottiger | IF (1:400) |
| PARP9 | Rabbit | Thermo (40-4400) | WB (1:500) |
| DTX3L | Rabbit | Bethyl (A300-834A) | WB (1:1000) |
| STAT1 phospho-Y701 | Rabbit | Cell Signalling (9167) | IF (1:400) and WB (1:1000) |
| FLAG | Mouse | Sigma (F1804) | IF (1:400) and WB (1:1000) |
| Anti-Rabbit-HRP | Donkey | Sigma (SAB3700934-2MG) | WB (1:5000) |
| Anti-Mouse-HRP | Donkey | Sigma (SAB3701105-2MG) | WB (1:5000) |
| Anti-Mouse-AF488 | Donkey | Thermo (A21202) | IF (1:400) |
| Anti-Rabbit-AF488 | Donkey | Thermo (A21206) | IF (1:400) |
| Anti-Mouse-AF568 | Donkey | Thermo (A10037) | IF (1:400) |
| Anti-Rabbit-AF568 | Donkey | Thermo (A10042) | IF (1:400) |
| Anti-Mouse-AF647 | Donkey | Thermo (A32787) | IF (1:400) |
| Anti-Rabbit-AF647 | Donkey | Thermo (A32795) | IF (1:400) |

## Acknowledgements

The authors wish to thank Anthony Fehr (Univ. Kansas) for the macrodomain vector used for cloning and fruitful discussions; Michael Hottiger (Univ. Zurich) for an ADP-ribose-binding reagents; Heli Alanen (Univ. Oulu) for production of the recombinant SARS-CoV-2 macrodomain; Célia Braga (Univ. São Paulo) for technical assistance; Priscilla Doria (Univ. São Paulo) for designing the graphical abstract and several colleagues from the University of São Paulo and from local pharmaceutical companies who helped source some of the compounds used for screening. The use of the facilities and expertise of the Proteomics and Protein Analysis core facility (a member of Biocenter Finland) and Biocenter Oulu sequencing center are gratefully acknowledged.

Work in the NCH lab is supported by a COVID19 emergency grant (2020/53017-6) and a Young Investigator Award (2018/18007-5) from FAPESP, LCR is funded by a FAPESP post-doctoral fellowship and held a Dimensions Sciences short-term fellowship, RT is funded by a PNPD/CAPES post-doctoral fellowship, ACM and IAM hold PhD scholarships from CNPq, the FCM lab is supported by a FAPESP Young Investigator II Award (2018/14898-2) and work in the ABC lab is funded by FAPESP grant 2019/26767-2. Work in the LL lab was funded by the Jane and Aatos Erkko Foundation and by the Academy of Finland (grant nos. 287063, 294085 and 319299 to LL).

## Author contributions

LCR and RT performed all of the cell-based experiments and RT and ACM performed all macrodomain cloning in the ABC and NCH laboratories. LCR and NCH developed the immunofluorescence analysis pipeline, IAM performed the virtual screen in the FCM laboratory, STS performed nanoDSF assays in the LL laboratory and both KD and NCH generated the CRISPR KO cells in the KWC laboratory. DS, FCM, ABC and NCH devised experiments and supervised the work, NCH managed the project and wrote the manuscript, with input from all authors.

---

**Supplementary Figure 1 – Image Analysis Pipeline**

1. Representative image of a complete sample acquire for a single well (DAPI channel only). Individual fields of view were stitched to form the shown image.
2. Representative image of a small portion of the sample, showing DAPI (blue) and ADP-ribose signal (red) in cells treated with IFNγ.
3. Representative image of the same cells as in 2, showing the ADP-ribose signal (grey) and the ring mask area in which ADP-ribose signal was quantified. Inner red circle is based on nuclei detection in the DAPI channel, outer red limit is either grown by 30 microns outwards from the nucleus (blue arrow), until neighbouring areas touch each other (yellow arrow) or until ADP-ribose signal reached background values (green arrow).
4. Scatterplots of DAPI area x Total DAPI intensity (right) or DAPI area x DAPI compactness (a measure of “roundness” of the nucleus) (left) for all nuclei detected in image 1. Each dot represents one nucleus, with clustering of nuclei represented by colour changes. Only nuclei within the shown gates were taken forward for quantification, as outliers were either incompletely acquired or not properly detected.
5. Representative image of the same cells as in 2, showing DAPI (blue), ADP-ribose signal (red), the ring mask area from step 3 (light green) and the detection of ADP-ribose dots (marked in yellow).
6. Representative column scatter plots of total ADP-ribose signal intensity contained in ADP-ribose dots per cell for three replicate samples performed in one experiment. Numbers of cells per sample are shown below each replicate. Each black dot represents one cell, red bars are mean + SEM.
7. Representative bar graph of ADP-ribose signal intensities. The mean signal intensity from 3000 to 5000 cells per replicate (red bar in graph 6) is plotted as a point, with the mean between replicates shown as a bar ± SEM.
8. For all figures in the manuscript, absolute signal intensities were normalized to the respective IFNγ-treated control (considered to be 100%), to normalize for variations in staining intensities between replicate experiments.

**Supplementary Figure 2 (related to Figure 2)**

**(A)** Representative image of immunoblot analysis for anti-FLAG and actin loading control in A549 cells transduced with empty vector control (e.v.) or with lentiviral constructs for constitutive expression of FLAG-tagged WT or N40A mutant macrodomain, 24h after treatment with vehicle control, 1000 U/mL IFNα, 1000 U/mL IFNβ, 100 U/mL IFNγ or transfected with 0.1 μg/mL poly(I:C).

**(B)** Example of the gating procedure used to normalize FLAG macrodomain expression levels. (left) Mean anti-FLAG immunofluorescence intensity in A549 cells transduced with empty vector control (e.v.) or with lentiviral constructs for constitutive expression of FLAG-tagged WT or N40A mutant macrodomain. (midle) Histograms of the FLAG signal intensity of the cell populations and gating strategy (red box) to select cells with comparable levels of macrodomain expression. (right) Mean FLAG immunofluorescence intensity in gated cells, showing comparable levels of expression of cells within this population, used for analyses in Main Figure 2C.

**(C)** Re-analysis of the results shown in Main Figure 2C using all cells, without the gating strategy described in Sup.Fig.2B.

**(D)** Representative images of the data shown in Main Figure 2C and Sup. Fig. 2C.

**(E)** Representative image of immunoblot analysis for anti-FLAG and actin loading control in A549 cells transduced with empty vector control (e.v.) or lenvitival constructs for doxyciline-inducible expression of FLAG-tagged WT or N40A mutant macrodomain, 24h after treatment with 100 U/mL IFNγ and indicated doses of doxycycline.

**(F)** Example of the gating procedure to normalize FLAG macrodomain expression between cell populations, similar to Sup.Fig. 2B above.

**(G)** Re-analysis of the results shown in Main Figure 2D using all cells, without the gating strategy shown in Sup.Fig.2F.

**Supplementary Figure 3 (related to Figure 3)**

**(A)** Quantification of ADP-ribose immunofluorescence signal intensity in A549 cells transduced either with empty vector control (e.v.) or with a lentiviral construct for constitutive expression of WT macrodomain, 24h after treatment with vehicle control, 100 U/mL IFNγ or 100 U/mL IFNγ + indicated doses of atorvastatin. Mean ± SEM (n=6, from 3 separate experiments). ****=p<0.0001.

**Supplementary Figure 4 (related to Figure 4)**

**(A)** Representative images of the results presented in Main Figures 4A and 4B.

**(B)** Representative image of immunoblot analysis for PARP9, DTX3L and actin loading control in RPE1-hTERT WT, PARP9 KO and DTX3L KO cells used in this study.

**(C)** Sanger sequencing traces obtained from PCR products of the genomic DNA surrounding the gRNA target locus in the PARP9 KO clone (top) and DTX3L KO clone (bottom) used in this study. The WT sequence and PAM location are shown above each trace for comparison, and the indel is highlighted with a red circle. Both clones are apparently homozygous for the shown allele, as the PCR product is expected to contain a mixture of amplicons from both alleles.

**(D)** Representative images of the results presented in Main Figure 4C.

**Supplementary Figure 5 (related to Figure 5)**

**(A)** Representative image (top) and quantification (bottom) of immunoblot analyses for STAT1-Y701 phosphorylation, FLAG and actin loading control in A549 cells transduced either with empty vector control (e.v.) or with a lentiviral construct for constitutive expression of WT macrodomain, 24h after treatment with vehicle control, 100 U/mL IFNγ or 0.1 μg/mL poly(I:C).

**Supplementary Tab1e 1.**
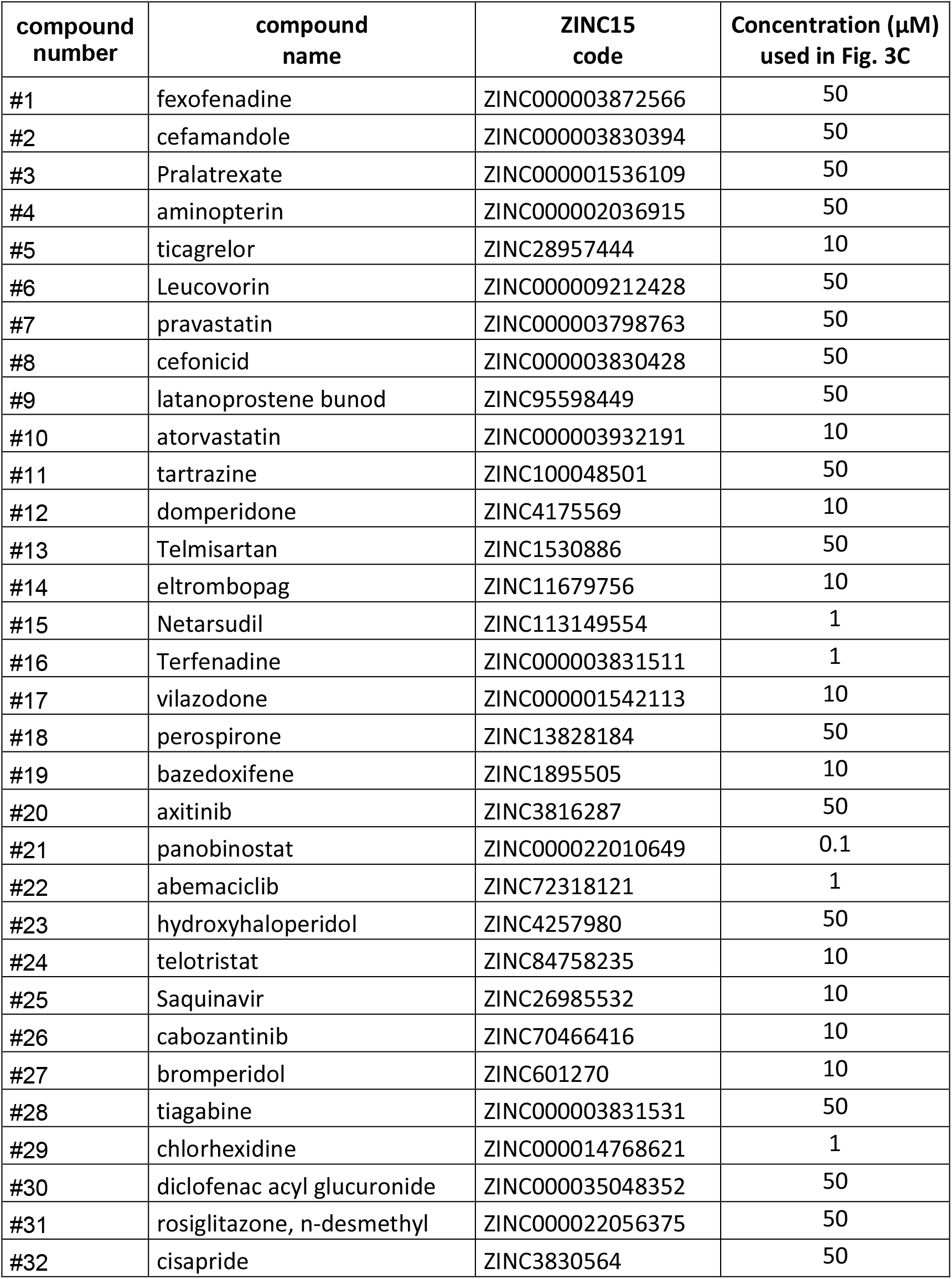
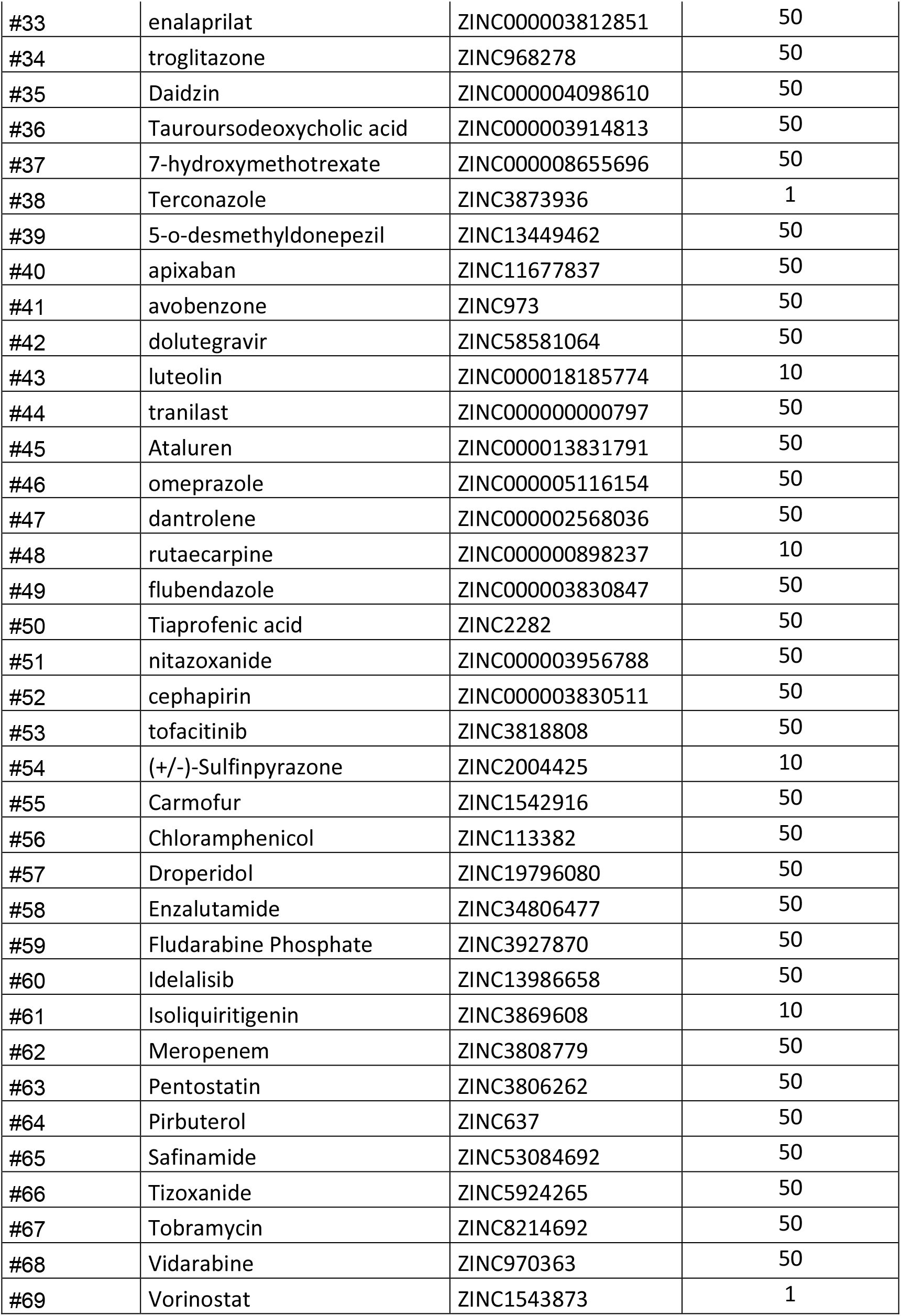
Compounds tested in the repurposing screen.

